# Plant-soil feedbacks trigger tannin evolution by niche construction: a spatial stoichiometric model

**DOI:** 10.1101/320572

**Authors:** Jean-François Arnoldi, Sylvain Coq, Sonia Kéfi, Sébastien Ibanez

## Abstract

Among plant traits, plant secondary metabolites such as tannins mediate plant-herbivore interactions but also have after-life effects on litter decomposition and nutrient cycling, which could influence their evolution. By modeling the flow of nitrogen and carbon through plants and soil in a spatially explicit context, we explored the relative contribution of herbivory and plant-soil feedbacks as drivers of tannin evolution. We assumed soil nitrogen to be composed of labile and recalcitrant compartments, the latter made of tannin-protein complexes accessible by plants via associations with mycorrhizal fungi. In unproductive environments and for plants with low biomass turnover rates, we show that when tannins modify soil properties locally, plant-soil feedbacks alone can drive their evolution. We further predict the existence of positive coevolutionary feedbacks between associations with mycorrhizae and tannins, possibly triggered by the evolution of the latter as protection against herbivores. In line with our theoretical results, tannins are mostly present in conservative plants, associated with mycorrhizae and inhabiting unproductive environments. Our work suggests that plant-soil feedbacks can enable niche construction mechanisms which can be key to the evolution of plant traits.

## Introduction

Through their impact on litter, soil and nutrient dynamics, plant traits are key drivers of ecosystem functioning. Many components of a plant’s environment are determined by the plant itself, each plant genotype shaping its own environment (Schweitzer et al., 2004; Whitham et al., 2006). In turn, these altered environmental conditions will affect the selective pressures acting on plant traits, thereby constituting an ecological-evolutionary feedback loop named *niche construction* (Lewontin, 2001; Odling-Smee et al., 2003). Many plant traits may have evolved because of their niche construction abilities, including flammability (Schwilk, 2003), germination timing (Donohue et al., 2005) and plant-soil feedbacks (Schweitzer et al., 2014). At the same time, environmental factors may interact with niche-constructing traits, either preventing or enhancing their evolution. For in-stance, flammability will more likely evolve in dry rather than wet environments. Most models of niche construction, however, do not include the interplay between environmental factors and niche constructing traits (Kylafis and Loreau, 2008; Laland et al., 1999; Lehmann, 2008; Schweitzer et al., 2014). Here, we propose that the evolution of tannin production in plants is a useful case study to investigate the evolutionary interplay between environment and niche constructing traits. Tannins can trigger plant-soil feedback loops because they complex organic nitrogen; and the importance of such feedbacks should depend on the fertility of the environment.

Among the large variety of plant’s defensive structures and compounds, tannins are a widely distributed class of plant secondary metabolites (Mole, 1993), with a wide array of intra-and interspecific variation in their production (Kraus et al., 2003). Although tannins have been extensively studied as anti-herbivore defense (Bergvall and Leimar, 2005; Feeny, 1970), they also affect nutrient cycling. After leaf senescence or plant death, they enter the soil as a component of litter, where they can greatly impact nutrient dynamics (Hättenschwiler and Vitousek, 2000; Joanisse et al., 2009; Kraus et al., 2003; Northup et al., 1995), limiting leaching (Jordan et al., 1979; Northup et al., 1995) and slowing-down decomposition and microbial activities (Benoit and Starkey, 1968; Coq et al., 2010; Joanisse et al., 2007; Schweitzer et al., 2004). Tannins complex macromolecules such as proteins which triggers the formation of recalcitrant organic matter, from which nutrient accessibility is reduced. This reduction of resource availability, however, may not only be a detrimental side-effect of an anti-herbivore strategy. Tannins promote local nutrient retention, which could be advantageous if plants can retrieve nutrients directly from recalcitrant organic matter, thereby circumventing mineralization by free microorganisms (Aerts and Chapin, 2000; Northup et al., 1995).

Mycorrhizal associations with symbiotic fungi may grant this ability to plants. Several studies have assessed the ability of the different groups of mycorrhizae to take up nutrients from organic matter (Read and Perez-Moreno, 2003), which was demonstrated for plants associated with ericoid mycorrhizae (Bending and Read, 1996; Joanisse et al., 2009; Wurzburger and Hendrick, 2009), which often abound in heathland and peatlands (we review results on the ability of different types of mycorrhizae to access recalcitrant organic matter in the Discussion).

For those mycorrhizal associations that permit N-accessibility from tannin-protein complexes, tannin production could therefore be a major process by which plants control their own resources, retaining nutrients in the local environment of the plant. This may affect the outcome of plant-microbe competition in favor of plants, and of tannin-rich *vs* tannin-less plants in favor of tannin-rich plants. This raises the question of the role plant-soil feedbacks in the evolution of tannins production, and the ecological conditions that could promote this role.

The observed correlation between nutrient limitation and tannin production (Endara and Coley, 2011; McKey et al., 1978) could therefore result from plants limiting nutrient losses through the formation of protein-tannin complexes, as a strategy to cope with an unfertile environment. Yet, one would expect the opposite correlation if the reduction of nutrient cycling by tannins is detrimental to plants (Vitousek, 1982). In fact, based on an eco-evolutionary model, and in apparent contradiction with empirical findings (Ordoñez et al., 2009), Barot et al. (2014) suggested that plants should evolve fast decomposing litter in nutrient-poor environments. Since tannins slow down mineralization, the question of whether the afterlife effects of tannins are beneficial or detrimental to plants in unfertile ecosystems is thus still open. If beneficial, tannin evolution could be driven by plant-soil feedbacks without additional selective pressure as an anti-herbivore defense. If detrimental, tannin evolution would be the outcome of a trade-off between anti-herbivore defense and the reduction of nutrient cycling.

Here we address this question and reconcile observations and theoretical predictions. To do so, we developed and analyzed a mechanistic evolutionary model, which includes the effects of tannins on both herbivory and plant-soil feedbacks. Because tannins are carbon (C) based compounds, and because their afterlife effect notably affects nitrogen (N) cycling, our model explicitly accounts for plants C:N stoichiometry. Because tannin-protein complexes are insoluble, immobile and remain near the tannin-producing plants, our model also accounts for spatial structure. Plants associations with specific mycorrhizae will be able to reabsorb N from tannin-protein complexes, in addition to labile N. Our model thus describes the ecological dynamics of plants growing in an N-limited substrate, plus the evolutionary dynamics of tannin production and symbiotic associations with mycorrhizae, in a stoichiometric and spatial context. We address the following questions:

1. Can plant-soil feedbacks trigger tannin evolution, even in the absence of herbivory?
2. What is the influence of ecosystem fertility on tannin evolution?
3. Can there be coevolution of symbiotic mycorrhizae and tannin production?

Our results suggest that for plants with low biomass turnover rates inhabiting un-fertile environments, spatial structure can promote the coevolution of tannin production and mycorrhizal associations. This can occur without selective pressure on tannins as a anti-herbivore defense, following a niche construction mechanism.

## Methods

Our results are based on the analysis of a model describing N and C fluxes through the plant-soil system. In the model, C is acquired by plants by photosynthesis, and N is acquired in the soil from either labile or recalcitrant N. Recalcitrant nitrogen is composed of tannin-protein complexes, and enters the soil system as litter produced by plants that produce tannins. Labile nitrogen can be absorbed by all plants. In contrast, recalcitrant nitrogen can only absorbed if plants are associated with mycorrhizae, which has a carbon cost. Finally, herbivores can remove biomass (and therefore C and N) from the plants. Herbivory is weaker when plants produce tannins. Two parameters are evolving: the proportion of tannins in plant biomass, *τ*, and the ability to associate with mycorrhizae that can use recalcitrant N, thereafter referred to as symbiotic capacity and denoted *e*.

We implemented the model at two levels of spatial complexity: a mean-field model assuming plants and soil to be well mixed in the landscape, and a spacially explicit model using a cellular automaton, with stochastic birth and death events.

### Meanfield model

#### Plant growth

Plant biomass is modeled by its amount of C and N content. We denote *B*_*τ*_ the C+N content of plant biomass, with C:N ratio *α*_*τ*_. A fraction *τ* of plant biomass is used to produce tannins, whereas the rest is involved in all other physiological processes, like growth, nutrient absorption or C exchange with mycorrhizae. Biomass is therefore constituted by a proportion *τB*_*τ*_ of tannin-C, and by the remaining biomass, *B* = (1 -*τ*)*B*_*τ*_, referred to as the effective biomass with C:N ratio *α*. We denote *γ* and *γ*_*τ*_ the proportion of N in *B* and *B*_*τ*_, respectively, with 1 -*γ* and 1 -*γ*_*τ*_, representing the complementary proportions^1^ of C in those compartments. At any given time, biomass growth is controlled by the limiting factor among C and N stocks (denoted *n* and *c*), i.e. the minimum between 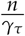 and 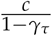 (Danger et al., 2008; Odum, 1971). Under these assumptions, and a turnover rate *h*, the dynamics of plant biomass *B*_*τ*_ follow

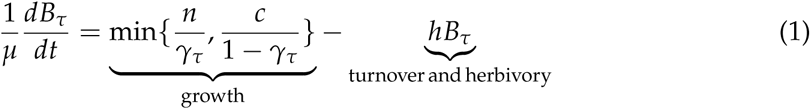

Here *μ* is a rate, which sets plants’ ecological timescale. We also model the effect of herbivory, that decreases plant biomass, as a part of the global turnover rate *h*. *h* is a decreasing function of *τ* when herbivores are present:

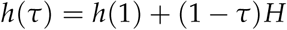

where (1 − *τ*)*H* is the relative rate of herbivory, reduced by tannins and reflecting the fact that they act as anti-herbivore compounds.

#### Resource acquisition by plants

In the soil, N is divided in two compartments: labile *n*_*l*_, and recalcitrant *n*_*r*_. Recalcitrant N is produced by plants themselves when tannins complex N-containing proteins during leaf senescence (Northup et al., 1995). Recalcitrant N is less subject to leaching and volatilization but is more difficult to absorb than labile N (Jordan et al., 1979). A critical assumption of our model is that symbiotic associations with mycorrhizae are necessary to absorb recalcitrant N (Pritsch and Garbaye, 2011; Read and Perez-Moreno, 2003; Wurzburger and Hendrick, 2009). We model this requirement by writing labile and recalcitrant absorption efficiencies (resp. *ℰ*_*l*_ and *ℰ*_*r*_) as explicit functions of *e*:

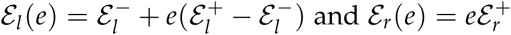

where 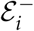 and 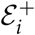 denote the extreme efficiency values. Thus, in our model, symbiotic capacity enhances the absorption capacity of both labile and recalcitrant nitrogen, but is necessary to absorb recalcitrant N. Being associated with mycorrhizae comes at a cost of maintenance *m*, linked to symbiotic capacity by

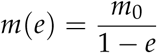

where *m*_0_ is the baseline maintenance cost. We note *ξ* the rate at which the absorption process is running, generating a cost *ξm*. On the other hand, plants access C by photosynthesis, at a rate *ϕ* which saturates, due to competition for light, as *B*_*τ*_ grows:

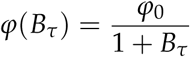

With this equation, *B*_*τ*_ is restricted to values smaller than one, thus defining a mass scale in our model. Under these assumptions, C and N stocks follow^2^, in physiological time scales *t*′, (which we will assume much shorter than the ecological time scale)

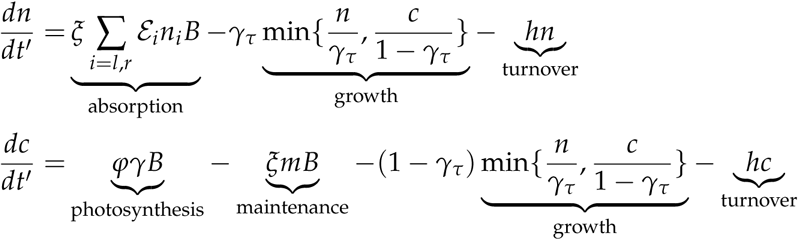

Physiological dynamics are best expressed in terms of concentrations *p* = *n*/*B* and *q* = *c*/*B* (Lemesle and Mailleret, 2008). Using 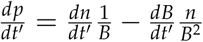 the above system becomes:

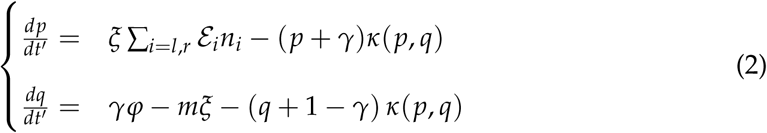

where 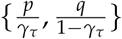. We assume physiological dynamics to rapidly reach a moving equilibrium (*p*_*_, *q*_*_) defined by 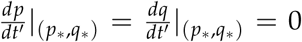. This equilibrium gives the plants stocks concentrations and depends on the pace of absorption *ξ*. We assume *ξ* to be plastic allowing plants to remain at stoichiometric balance, implying that *p*_*\**_/*q*_*\**_ = *α*_*τ*_. This assumption and eq. (2) together fix the value of the absorption rate *ξ* ∑_*i*=*l,r*_ *ε*_*i*_*n*_*i*_. It will be convenient to write the latter as *γ𝒜*. With this notation:

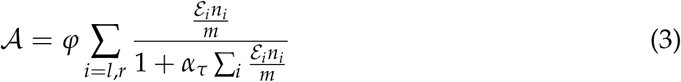

This expression proposes a natural decomposition *𝒜* = *𝒜*_*l*_ + *𝒜*_*r*_, representing the sum of absorption rate of labile and recalcitrant N, respectively. Stoichiometric balance also implies that *κ*(*p*_*\**_, *q*_*\**_) = *p*_*\**_/*γ*. We define *P* = *p*_*\**_/*γ*, which corresponds to the ratio of N-stocks over the N-content of effective biomass, *γB*. From eqs. (2-3) it holds that

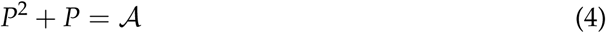

Eq. (1) defining plant growth can finally be rewritten:

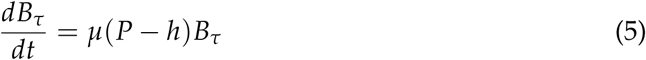

Productivity (rate of biomass creation) equals *µP*, therefore *P* is the effective nutrient stock which drives plant productivity in our model. That *P* satisfies Eq. (4) implies that productivity follows a type-II functional response to soil N content.

#### Nutrient absorption and recycling

In the soil, we model two variables: the amounts of labile and recalcitrant nutrients (*n*_*l*_ and *n*_*r*_, respectively). The dynamics of labile nutrients is controlled by three fluxes: absorption by plants, input by litter fall, and exogenous nutrient flows.

Absorption by plants is modeled as described in the previous section on plant physiological dynamics removing N from the soil at a rate *µ𝒜*_*l*_ *× γB*. Litter input depends (i) on the amount of litter entering the soil and (ii) on the proportion of litter N that is in a labile form. The amount of litter entering the soil is proportional to turnover rate and the N parts of biomass, *γB*, plus that of stocks, *n* = *PγB*. The proportion in labile form is a negative function of plant tannins, because tannins will complex part of nitrogen and turn it into its recalcitrant form. If one unit of tannins complexes Ω units of N, then we get a recycling term *µ ℛ*_*l*_ *× γB*, with

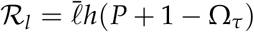

where 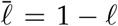 models N losses during decomposition (e.g. volatilization) and Ω_*τ*_ = Ω*τ*/*γ*_*τ*_ so that Ω_*τ*_*γB* is the amount of N biomass being complexed by tannins. The dynamics of labile soil nitrogen are completed by an exogenous flow, representing natural input and output of nitrogen, in particular through leaching. We write this term as

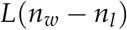

Where *n*_*w*_ is the bare soil nutrient content (Droop, 1968; Lemesle and Mailleret, 2008) and *L* is the leaching rate of labile N. We will take this rate to set a reference ecological time scale, so that henceforth *L* = 1. As a result, the global equation modeling the dynamics of labile N in the soil is:

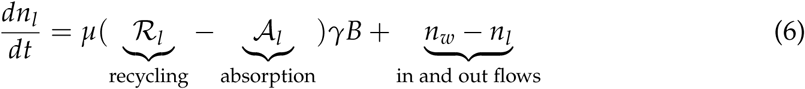

What N is complexed by tannins flows into the recalcitrant compartment *n*_*r*_. This is the only inflow in *n*_*r*_. Similarly as the one of labile N, the dynamics of *n*_*r*_ thus read

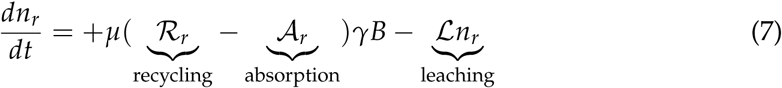

*ℛ*_*r*_ = Ω_*τ*_. Note that symbiotic capacity *e* only enters in the absorption terms, via the benefit to ratio *ε*_*i*_/*m*. The mean-field model is determined by the system of ODEs (5-6-7) complemented by eqs. 3 and 4.

#### The cellular automaton and the inclusion of explicit space

Because plant-soil feedbacks act at a local scale, we implemented a spatialized model using a cellular automaton. Space was represented as a landscape of S patches, which can be either empty or occupied by an individual plant.

Individuals grow according to the above model, absorbing and recycling N locally. But, in contrast to the mean-field model, N diffuses continuously across patches. For a general topology of connected patches the diffusion term affecting nutrients in patch *k*, either labile 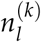 or recalcitrant 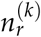, reads,

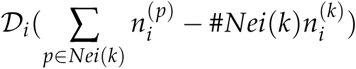

where *Nei*(*k*) denotes the set of neighbors of patch *k* and *𝒟*_*i*_, *i* = *l, r*, are diffusion rates. The first term (the sum) represents in-flow from neighboring patches of patch *k* while the second represents out-flows from patch *k* towards its neighbors. In simulations we considered the simplest possible landscape: a one-dimensional periodic lattice, so that *Nei*(*k*) = {*k -*1, *k* + 1} mod(*S*). We will typically assume *𝒟*_*l*_ to be much larger than *𝒟*_*r*_, making recalcitrant N less mobile. We then modeled population dynamics. This was implemented by allowing individuals to randomly die and/or reproduce in an empty patch. We assume that larger individuals are less likely to die than smaller ones and, similarly, that production of seedlings increases with biomass. Concretely, we set

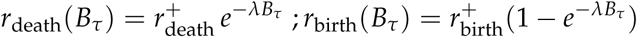

Newly produced plants colonize empty patches. The probability of establishment of a seedling is higher in patches located near the parent tree and lower for distant patches.

Concretely, we write the recruitment rate in an empty patch *k* as

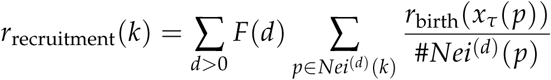

where *F*(*d*) is the fraction of a plant reproductive effort that will result in colonizing patches located at distance *d* from this parent plant (*Nei*^(*d*)^(*k*) is the set of patches at distance *d* from patch *k*). In simulations we will superpose a geometrically decreasing kernel up to *d* = 3 to a uniform contribution of the remaining effort to all other patches, representing the fraction of seeds transported by the wind or by animals to arbitrarily distant locations. Concretely, for some *δ, K <* 1, *F*(1) = (1 *-δ*)*K, F*(2) = (1 *-δ*)*K*(1 *-K*), *F*(3) = (1 *-δ*)(1 *-K*)^2^ to which we add a mean-field contribution 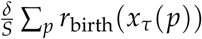.

When a new plant is recruited in a patch, the plant starts growing from a small initial biomass. Generally, the demographic parameters were tuned to always have a densely populated landscape. Indeed, if many patches are empty at any given time, the dynamics become neutral and we cannot expect to exhibit selective pressures (see Table 1).

**Table 1:** Model parameters (in order of appearance). Rates are expressed relatively to labile leaching rate. Similarly, the biomass density of plants that saturate photosynthesis sets the mass scale. Recall that the other stoichiometric parameters *γ, α*_*τ*_ and *γ*_*τ*_ are explicit functions of *α* and *τ*.

| Parameter | interpretation | value |
| --- | --- | --- |
| $\alpha$ | C:N ratio of effective biomass | 0.5 |
| $\mu$ | metabolic rate | 5 |
| $h$ | relative turnover rate | 0.6 |
| $H$ | relative herbivory rate | 0 or 0.01 |
| $\mathcal{E}_l^-$ | min labile absorption efficiency | 1 |
| $\mathcal{E}_l^+$ | max labile absorption efficiency | 2 or 2.1 |
| $\mathcal{E}_r^+$ | max recalcitrant absorption efficiency | 1.1 |
| $m_0$ | min maintenance cost | 1 |
| $\varphi_0$ | max photosynthesis rate | 10 |
| $\ell$ | N volatilization | 0.2 |
| $n_w$ | bare soil N content | 1 |
| $\mathcal{N}$ | effective bare soil N content | 0.6 |
| $\mathcal{L}$ | recalcitrant leaching rate | 0.1 |
| $\Omega$ | Tannins complexation power | 1 |
| $\mathcal{D}_l$ | labile diffusion rate | 10 |
| $\mathcal{D}_r$ | recalcitrant diffusion rate | 0.1 |
| $r_{\text{birth}}^+$ | max birth rate | tuned to |
| $r_{\text{death}}^+$ | max death rate | avoid |
| $\lambda$ | demographic effect of biomass | neutrality |
| $K$ | seed dispersal kernel | 0.6 |
| $\delta$ | fraction of seeds uniformly dispersed | 0.2 |
| $z$ | phenomenological local mass parameter | 0 or 0.03 |

## Model analyses

### Invasibility of the tannin phenotype

We study the conditions for tannin producing plants to invade a resident population of tannin-less plants. We suppose for now that all plants can absorb recalcitrant N, with equal and constant symbiotic capacity (this assumption will be relaxed for the study of tannin - mycorhize coevolution). In simulations we will assume that *ε*_*l*_ = *ε*_*r*_ but our analysis allows for any choice of non zero efficiencies. We focus on the influence of two key parameters, the effective bare soil N content and the plants’ relative turnover rate *h*.

The effective bare soil N content, *𝒩*, is a nondimensional combination of parameters:

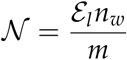

It is a proxy for soil fertility, as perceived by plants, encompassing any choice of efficiencies *ε*_*i*_ and maintenance function *m*. Changing *𝒩* will allow us to test if the model predicts that unfertile ecosystems favor tannin production, as observed in nature. Plants relative turnover *h* is a proxy for plant life strategy, high *h* plants will enhance nutrient cycling while low *h* plants will slow it down, relative to their growth (here controlled by the rate *µ*). Our reasoning relies on the analysis of the invasion fitness function (Geritz et al., 1997), i.e. the initial growth rate of a mutant population in the presence of a resident population at equilibrium. In some simulations we will directly access the outcome of an invasion attempt, but the basic idea is the same. A resident population sets the state of soil labile N and available light that a rare mutant population *B*_*τ*_ will experience. The mutant’s fitness reads

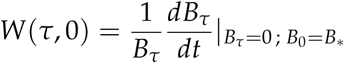

where by definition *W*(0, 0) = 0. Its gradient *∂*_*τ*_*W*(*τ*, 0) determines invasibility: if positive, a mutation leading to a small change in the phenotype *τ* can invade and the phenotype is consequently selected. In our model:

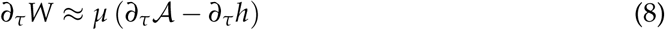

If the phenotypic change increases the absorption rate *𝒜* and/or decreases the turnover rate *h*, mutants will invade. In our model, tannins can be selected as a protection against herbivores because they decrease turnover (*∂*_*τ*_*h <* 0).

In the absence of herbivores the selective advantage of tannins can only come through an increase of absorption (*∂*_*τ*_*𝒜 >* 0). However, an increase of absorption would imply that mutants, by producing tannins, have transformed the soil composition to their advantage, so that the cost of tannin production is worth the benefits of an increase of the recalcitrant nutrient pool. Because mutants are rare by definition, this cannot happen if they are well mixed in the landscape: locally, the recalcitrant pool would be negligible. However, spatial structure may allow mutants to modify their local N-environment.

In a first step we will assume that mutants, via some spatial mechanisms such as aggregation, *do* modify they local N-environment. We call this approach spatially implicit, as it does not explicitly address the origins of the local modification of the resource environment by mutants, but assume this modification to take place (we detail this reasoning below). Using this approach, we consider three evolutionary scenarios, the first setting a baseline by neglecting spatial structure:

i. Tannin phenotype evolving in the presence of herbivores, without taking into account spatial structure.
ii. Tannin phenotype evolving without herbivory pressure but implicitly taking into account spatial structure.
iii. Tannin phenotype evolving in the presence of herbivores, implicitly taking into account spatial structure.

In a second step we will test the predictions of the first step using simulations of the cellular automaton. We will then use the cellular automaton to analyze the “social” effects allowed by spatial structure, such as altruism (van Baalen and Rand, 1998) and ecological inheritance (Odling-Smee et al., 2003). In other words, after analyzing the consequences of a local modification of the environment by tannins (as in the implicit approach), we explored the underlying spatial mechanisms.

#### First step: implicit spatial structure

Suppose that tannins affect the soil composition surrounding one or a clustered group of mutants. We account for this by defining a parameter *z*, an implicit function of spatial processes controlling aggregation of mutants and the diffusion of the recalcitrant N that they produce. To define *z*, we start from 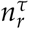, the amount of recalcitrant N formed in the vicinity of a mutant individual, or group, with biomass *B*_mut_. This N content should be proportional to *τB*_mut_, the C-amount of tannins produced by the mutants. This will indeed induce an inflow into the local pool 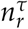 proportional to *τB*_mut_ *× μh*, where *μh* is the absolute biomass turnover rate. At the same time, this pool will suffer a loss at a rate proportional to leaching and diffusion. Keeping track of relative turnover *h*, and tannins *τ*, we define *z* so that 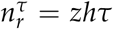. This phenomenological mass parameter will enable us to tune the direct effect that mutants, by changing their N-environment, have on their absorption effort *𝒜*. Concretely we evaluate eq. 8 at *τ* = 0, with a change in absorption computed as

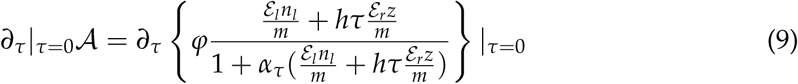

and evaluated at the equilibrium values of the resident. Note that setting *z* = 0 corresponds to the mean-field model in which there is no spatial structure.

This analysis answers the question: – Given that mutants modify their local environment (as measured by *z*), under which conditions will this allow them to invade a resident population, and thus lead to the selection of the tannin phenotype?

#### Second step: explicit spatial structure

We used the cellular automaton to test the results of the implicit approach described above. We simulated invasion attempts of tannin-producing plants (with *τ* = 0.2) in a landscape populated by tannin-less residents. The landscape was a periodic lattice comprised of *S* = 250 cells. Invasions started from 5 spatially aggregated individuals. We monitored the ultimate fraction of mutants after 400 time steps. A snapshot of a run of the automaton is presented in Fig 2, where we can see the impact of tannin producing individuals on soil variables surrounding their respective patches.

**Figure 1:**
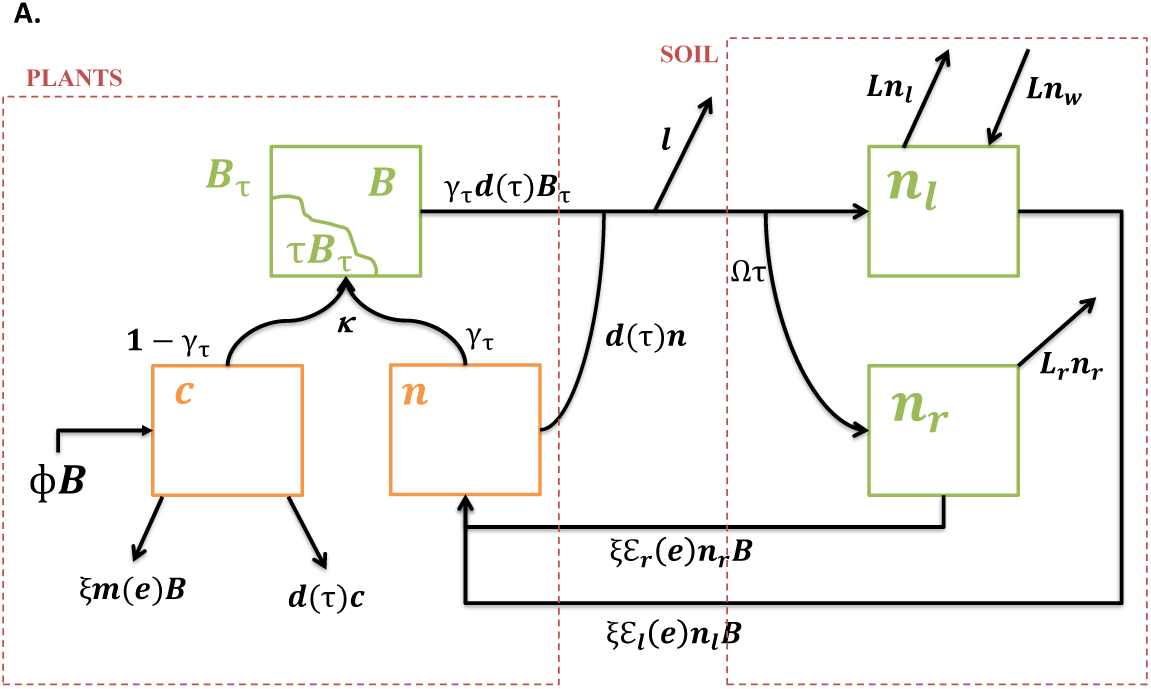
Scheme of the mean-field model. On the left, the three plant compartments. Carbon (*c*) and nutrient (*n*) stocks are used by the plant for the growth of its effective biomass (*B*_*τ*_), which includes tannins. Following senescence, nutrients enter one of the two soil compartments, on the right. Nutrients complexed by tannins enter the recalcitrant nutrient pool (*n*_*r*_), the remaining nutrients enter the labile nutrient pool (*n*_*l*_). Nutrients are then absorbed by plants, at the expense of carbon costs.

**Figure 2:**
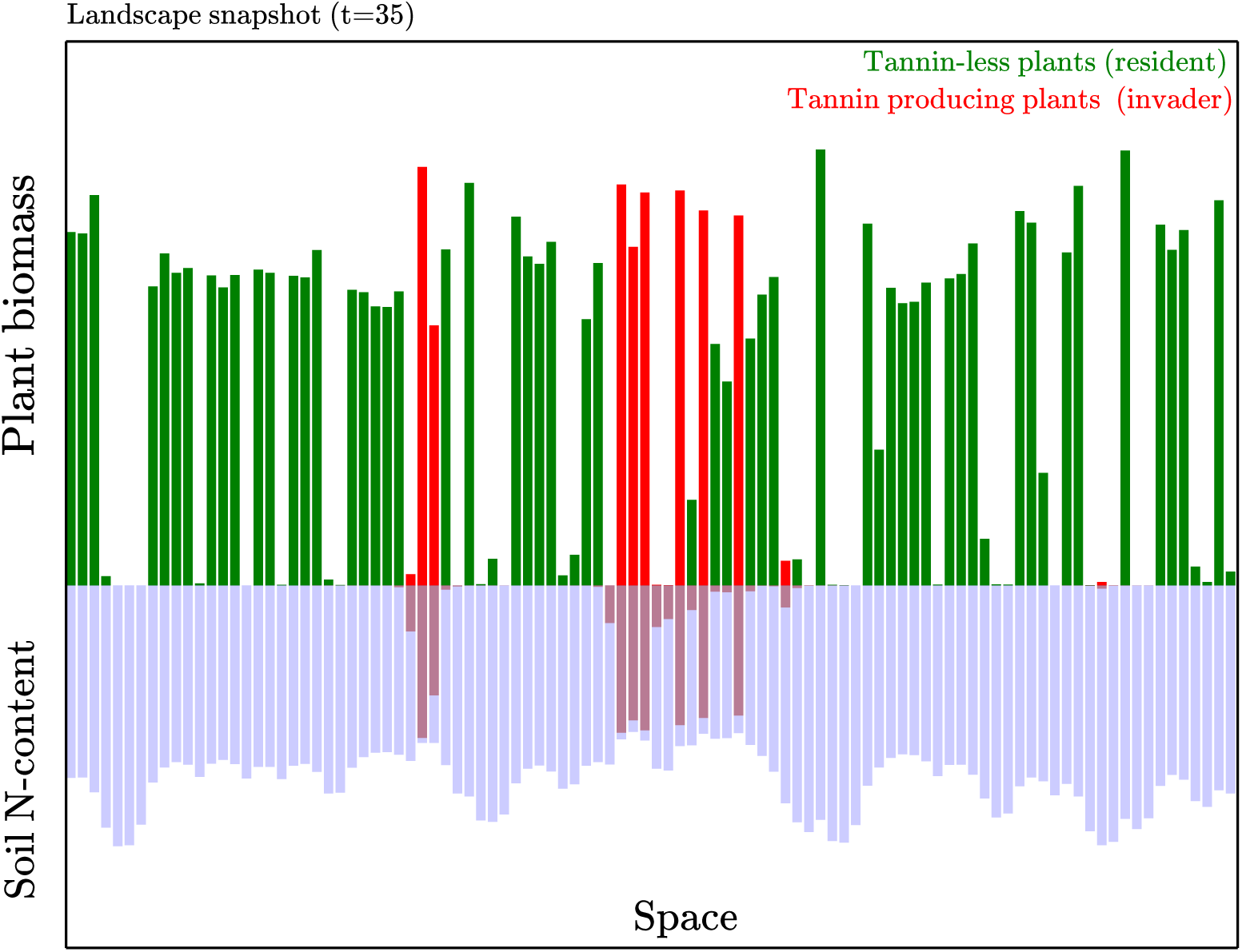
The cellular automaton (snapshot). Each green or red bar represent the biomass of an individual plant. Inverted bars represent soil N-content in its labile and recalcitrant form (blue and brown respectively). We follow here the invasion attempt of tannin producing plants (in red) in a landscape populated by a tannin-less resident population (in green).

We also used the cellular automaton to focus on social effects allowed by spatial structure. 1) To relate tannin evolution to the evolution of altruism in a spatial context (van Baalen and Rand, 1998), we ran invasion attempts along a gradient of values of recalcitrant diffusion rate *𝒟*_*r*_. We deduced the probability of a successful invasion as a function of *𝒟*_*r*_. The idea was that, for low diffusion, mutant individuals are effectively “selfish”: they do no share their complexed N-pool with neighboring plants. By contrast they share it with all plants if diffusion is large. In this latter case we expect a “tragedy of the commons”, where the resident plants, seen as “cheaters”, benefit without costs from the complexed N pool so that tannins are not selected. For intermediate diffusion, a tannin producing individual shares his complexed N pool with a few other individuals that are likely to be of his kin. If invasion is more likely then at *𝒟*_*r*_ = 0, this would demonstrate the contribution of group selection, a mechanism that can allow the selection of pure altruism (van Baalen and Rand, 1998; Wilson, 1980). 2) The death of a tannin producing individual leaves a N-rich patch available for recruitment from neighbors that are also likely to be of his kin. In this case individuals inherit the environment transformed by their ancestors, a property termed ecological inheritance (Odling-Smee et al., 2003). To test if ecological inheritance played a role in our system, we devised an arbitrary way to turn it off: following the death of a tannin producing plant, we set to zero the recalcitrant N pool that it had created (i.e. N pools vanish before the establishment of new seedlings). This requires diffusion of recalcitrant N to be negligible, otherwise it becomes impossible to determine the part of *n*_*r*_ that is associated to the individual plant that died. Thus, setting *𝒟*_*r*_ = 0 we monitored the ultimate relative abundance of mutants, with and without ecological inheritance.

### Coevolution of tannins and symbiotic capacity

For any resident phenotype (*e, τ*), we computed the ecological equilibrium of the mean-field model. We then determined the fitness gradient at (*e, τ*) by evaluating the initial growth rate of slightly different mutant phenotypes (*e* + *δe, τ* + *δτ*). The fitness gradient has now two components, one in the direction of symbiotic capacity and one in the direction of tannins:

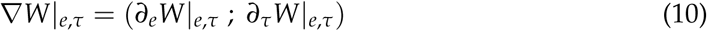

Coevolutionary trajectories follow the direction of the fitness gradient until it vanishes and a steady state is reached (Leimar, 2009). For the evolution of single traits, such steady states are called CCS, Continuously Stable Strategies (Eshel, 1983; Geritz et al., 1998); in a coevolutionary context we will refer to co-CSS.

Similarly as in eq. 8 from the invasibility analysis, the first component of the fitness gradient is proportional to *∂*_*e*_*𝒜*, the change of N-absorption as *e* is varied. In our model, this change is driven by nondimensional ratios representing the effective fertility of the soil as perceived by the plants, which read:

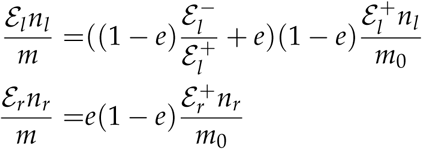

They are non monotonous functions of *e*, meaning that there is a trade-off between increasing efficiency and reducing C-cost.

The second component of the fitness gradient is proportional to *∂*_*τ*_*𝒜 − ∂*_*τ*_*h*, where *∂*_*τ*_*h* is non zero only in the presence of herbivores. The absorption term depends, in particular, on the way mutants change their local N-environment. To model the outcome of this change, we used eq. 9 which implicitly accounted for spatial structure via the phenomenological mass parameter *z*. We slightly generalized the approach to allow for a tannin producing resident. We simply wrote the recalcitrant N-content in the vicinity of an aggregated group of mutants as

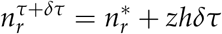

where 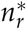 is recalcitrant content produced by the tannins of the resident plants. From there we applied the same reasoning as in eq. 9 to compute *∂*_*τ*_*𝒜*.

The co-CSS will correspond to an intersection between the isocline of symbiotic capacity {(*e, τ*) *| ∂*_*e*_*W|*_*e,τ*_ = 0} and the one of tannin content {(*e, τ*) *| ∂*_*τ*_*W|*_*e,τ*_ = 0}. We first chose 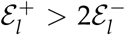 so that symbiotic capacity would be favored in the absence of tannins by enhancing labile absorption. We asked whether this could then initiate the evolution of tannins, and if so, if this would further promote symbiotic capacity. By contrast, we considered the case where herbivory could favor tannin production but where symbiotic capacity could not evolve without tannins (i.e. 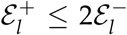). We asked whether the evolution of tannins as an anti-herbivore defense could then promote the evolution of symbiotic capacity and then further promote tannin production.

## Results

### The spatially implicit approach predicts that unfertile environment and intermediate turnover favor tannin evolution

The relationship between tannin invasibility, soil fertility and plants life strategies under the three scenarios is illustrated in Fig. 3, in which we numerically computed the fitness gradient (eq. 8) as a function of *h* and *𝒩*. In the leftmost panel – herbivores and no spatial structure – we observe selective pressure towards protection against herbivores by tannin production at low turnover rates *h* and relatively low fertility *𝒩*. Note that as *h* grows (or *𝒩* decreases) plants eventually reach the limits of viability (in gray in Fig 3). In the middle panel – no herbivores but spatial structure – we observe that local recycling of recalcitrant N by tannin-producing mutants should strongly promote their settlement at low fertility and intermediate values of turnover rate. Finally, the rightmost – herbivores and spatial structure – combines the previously described behaviors, showing that, for low soil fertility, tannin production can be beneficial at both low and intermediate values of turn over rates, but for essentially different reasons. At low turn over rate tannins are mostly selected for their herbivore deterrent properties, whereas at intermediate turnover they mostly favor plants by forming a local N pool less subject to leaching.

**Figure 3:**
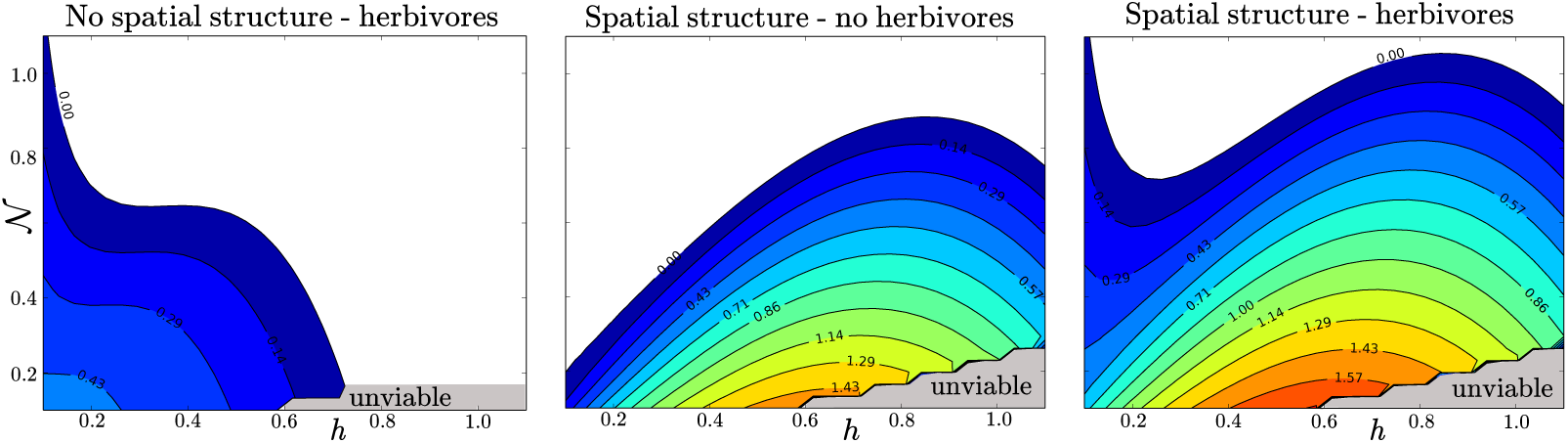
Role of turnover rate *h* and effective soil fertility *𝒩* on the invasibility of tannins. We plotted the fitness gradient at *τ* = 0, a positive value predicts that tannin-producing plants should be selected for. In all panels, the facts that the limits of viability (shaded gray) increases in steps is a numerical artifact. Left panel: non spatial model (*z* = 0) with herbivory (*H* = 10^*-*2^). In this case, recalcitrant N play no role in the invasion fitness of tannins. middle panel: Spatially implicit context without herbivory (*z* = 0.03). Right panel: spatial structure and herbivory.

### Simulations of the cellular automaton confirm the predictions of the implicit approach and demonstrate the importance of social effects

In Fig. 4 we confronted the qualitative predictions of the implicit approach to simulations of the cellular automaton, for scenario (ii) i.e. no herbivory. For the same values of parameters as in Fig 3 we ran simulations of invasions attempts of a tannin producing population (*τ* = 0.2), starting from a few aggregated individuals in a landscape populated by tannin-less plants. We monitored the ultimate fraction of mutants and compared the outcome of these simulations to the invasibility domain from the middle panel of Fig 3. Except near the inviable domain where less successful invasions were recorded than expected, we see that simulations of the cellular automaton are in qualitative agreement with the invasibility predictions.

**Figure 4:**
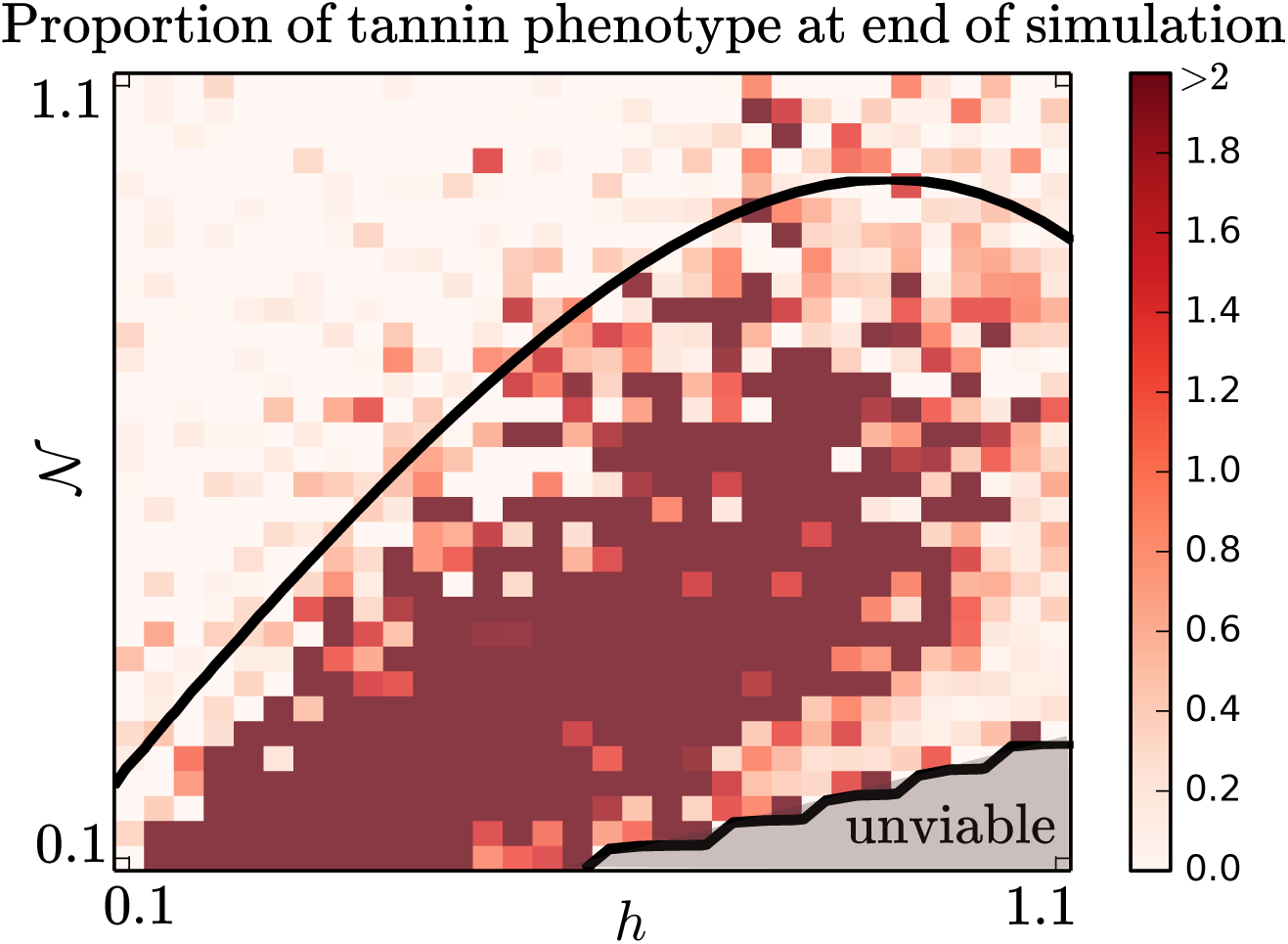
Confrontation of the qualitative predictions of fig.(3) to simulations of the cellular automaton. Each cell represents the ultimate fraction of mutants (with *τ* = 0.2) after 400 time steps. The landscape is a periodic lattice comprised of *S* = 250 cells. We start from 5 spatially aggregated mutants in a landscape populated by tannin-less plants. New-born individuals have an initial biomass 10^*-*3^ smaller than the one of an average resident. N-diffusion rates are set to *𝒟*_*l*_ = 5 and *𝒟*_*r*_ = 0.1. The invasibility domain from the middle panel of Fig. 3 is reported as a black line (*z* = 0.03). Except near the inviable domain where less successful invasions are observed than expected, we observe a good qualitative agreement between the prediction of the implicit approach and simulations.

Beyond corroborating the analysis from the implicit approach, simulations of the cellular automaton allowed us to relate tannin evolution to the evolution of altruism by group selection (van Baalen and Rand, 1998). In the left panel of Fig 5 we plotted the probability of a successful invasion as a function of recalcitrant diffusion rate *𝒟*_*r*_. At the individual level, in terms of survival and birth rate, the least diffusion of recalcitrant N the better *-*a selfish “behavior”. Yet at the group level, in terms of probability of invasion, we observe that intermediate values of diffusion – altruistic behaviors – aremore beneficial. For larger values of nutrient diffusion “cheaters” benefit without costs from the complexed N pool, and individuals contributing to this pool, having no extra benefits, are eventually excluded.

**Figure 5:**
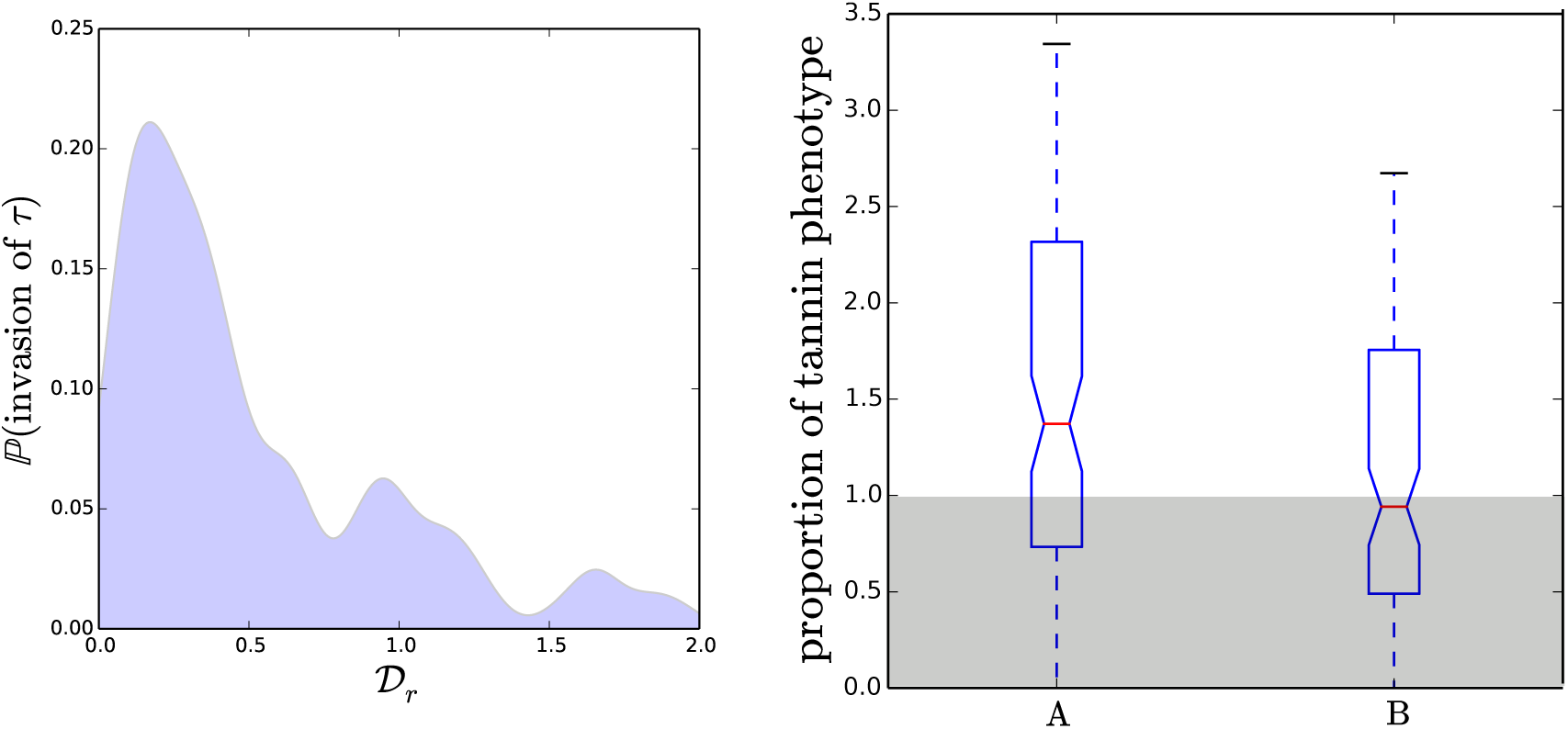
Social effects promoting tannins (simulations of the cellular automaton) Left: Altruism. We varied the recalcitrant diffusion rate *𝒟*_*r*_ from 0 to 2 and estimated the invasion probability function from the histogram of diffusion rates for which the ultimate proportion of mutant biomass was larger than one. At the individual level, in terms of survival and birth rate, the least diffusion of recalcitrant N the better. Yet at the population level, in terms of probability of invasion, intermediate values of diffusion appear more beneficial. For strong diffusion we recover a “tragedy of the commons” scenario: “cheaters” benefit without costs from complexed N. Right: ecological inheritance. A: the box plot represents the outcome of simulation runs (100 runs up to t=200), starting from a few tannin producing-mutants in a landscape populated by a tannin-less resident. B: when an individual died, the recalcitrant pool that it formed during its lifetime is set to zero, thus removing ecological inheritance. The comparison between case A and B demonstrates a positive effect of ecological inheritance on the selection of tannins.

The death of a tannin producing individual leaves a N-rich patch available for recruitment from neighbors that are likely to be of his kin, a property termed ecological inheritance (Odling-Smee et al., 2003). As illustrated in Fig 5, this effect was positive in our model: without ecological inheritance (case B in Fig 5) the invasion of tannin producing mutants is substantially slower.

### Positive coevolutionary feedbacks between tannins and symbiotic capacity can be triggered by herbivores

In Fig. 6, using the implicit account of spatial structure, we could sketch the coevolutionary dynamics in trait space (*e, τ*), generated by the fitness gradient eq. 10 (Leimar, 2009). The fitness gradient has two components and the co-CSS corresponds to an intersection between the fitness gradient isocline of symbiotic capacity (in gray in Fig. 6) and the one of tannin content (in black).

**Figure 6:**
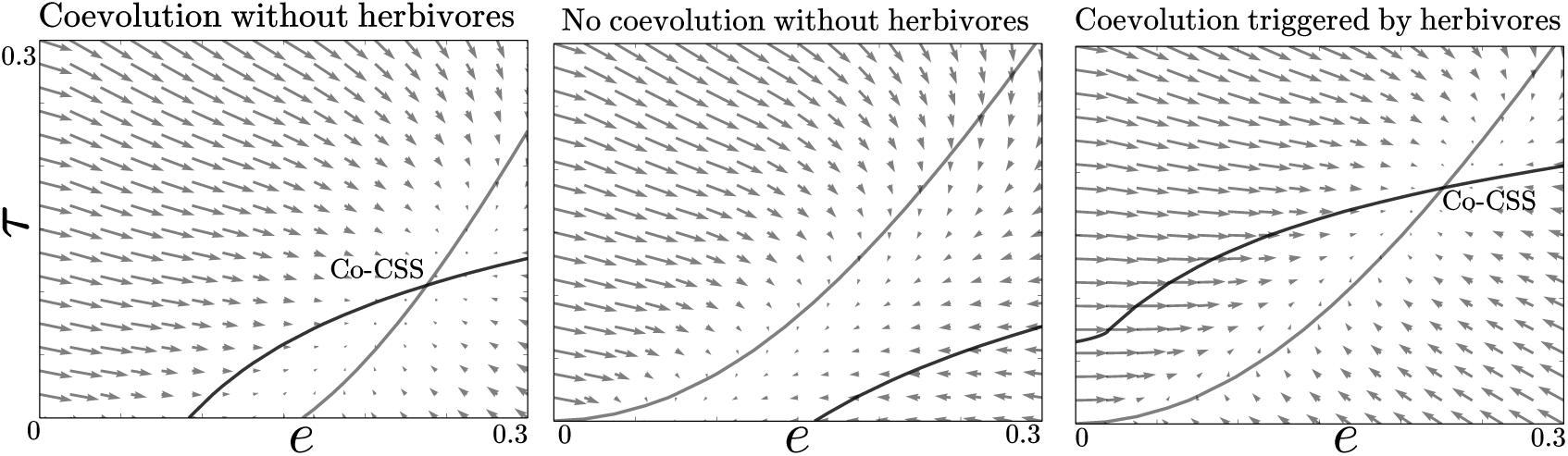
Coevolution of tannins and symbiotic capacity. In gray is the fitness gradient isocline of symbiotic capacity, In black the fitness gradient isocline of the tannin phenotype. The coevolutionary Continously Stable Strategy (co-CSS) is at the intersection of the two. Leftmost panel: we chose parameters of absorption efficiencies so that that symbiotic capacity would be favored even in the absence of tannins. In this case, we predict a coevolutionary positive feedback, with tannin evolution increasing the selective advantage of symbiotic associations and vice versa, until a co-CSS is reached. Middle panel: here we chose absorption efficiencies so that symbiotic capacity is not favored in the absence of tannins. In this case, we predict no evolution of tannin production nor symbiotic capacity. Rightmost panel: Additional pressure from herbivory, however, can trigger a coevolutionary positive feedback.

In the first panel, labile absorption is suficienltly inefficient that symbiotic capacity can be selected even in the absence of recalcitrant N (the fitness gradient along *e* is positive at *e, τ* = 0). We observe a positive coevolutionary feedback: if symbiotic capacity increases, complexed N can be absorbed so that tannins can become beneficial, which in turn enhances the selective pressure on symbiotic capacity, and so on until the co-CSS is reached.

In the middle panel, symbiotic capacity is too costly to evolve in the absence of tan-nins, and no coevolution is possible. However, in the rightmost panel, herbivory allows the evolution of tannins as an anti-herbivore defense. This triggers a positive coevolutionary feedback: symbiotic capacity evolves which can enhance tannins until the co-CSS is reached. This demonstrates the potential complementary of the two hypothesis concerning tannin evolution in plants (i.e. anti-herbivore defense and niche construction): the fact that tannins form anti-herbivore compounds can be at the origin of the plant-soil feedbacks that can, in turn, promote tannin production and mycorrhizal associations.

## Discussion

### Plant soil feedbacks alone can trigger tannin evolution

Our results are in line with the “litter perspective” proposed by Hättenschwiler et al. (2010) as an alternative to the “green foliage perspective” for the evolution of tannins. Plants benefit from the afterlife effects of tannins, instead of suffering from a reduction of nutrient cycling (Barot et al., 2014). In line with empirical findings (Joanisse et al., 2009; Kraus et al., 2004), our model assumes that the main mechanism explaining the impact of tannins on the soil is their ability to complex proteins during leaf senescence, thus forming a recalcitrant N pool that can be reabsorbed by mycorrhizae.

This can confer a selective advantage to tannin-producing plants with intermediate biomass turnover rates (Figs 3–4). If turnover is too slow, plant biomass accumulates, photosynthesis becomes limited by intraspecific competition and the carbon stock is to low for plants to invest in tannins. By contrast, if turnover is too fast, labile nutrients abound and the recalcitrant pool formed by tannins becomes useless. Importantly, tannin-producing plants must have a preferential access to the recalcitrant pool. The mechanisms underlying such exclusivity may be spatial (as in our model), or biological, if mycorrhizae are better adapted to reabsorb the recalcitrant nutrients formed by their host (Joanisse et al., 2009; Wurzburger and Hendrick, 2009). Tannin production cannot evolve when the recalcitrant pool is shared by all plants, because tannin producing plants must be protected from the invasion of cheaters able to absorb recalcitrant N without paying the cost of tannin production.

Our model is in this sense similar to those focused on the evolution of altruism (Le Galliard et al., 2003), facilitation (Kéfi et al., 2008) and shared chemical resources among microorganisms (Allen et al., 2013). Yet our model is not a mere illustration of altruism evolution. Indeed, in models of altruism evolution (Le Galliard et al., 2003; van Baalen and Rand, 1998; Wilson, 1980), the altruistic trait is beneficial to any phenotype. Our model is one of competition for resources in which no individual is strictly altruistic. In principle, tannin production could both lead to the competitive exclusion of non-symbiotic plants (Joanisse et al., 2009) and to the facilitation of plants associated with mycorrhizae. Finally, we considered here a whole ecosystem including abiotic compartments, instead of a single monospecific population. Hence, selection does not operate at the group-level (van Baalen and Rand, 1998), but at the ecosystem-level (see *Tannin evolution under a niche-construction perspective* below).

### Nutrient-poor ecosystems favor tannin evolution

Because they slow down the mineralization of recycled nutrients, one could have expected tannins to be detrimental to plants in unfertile ecosystems (Barot et al., 2014). This prediction, however, is not verified empirically, both at the inter and intraspecific level: tannin producing species are mostly found in unfertile environments (Endara and Coley, 2011; McKey et al., 1978), and plant populations of a single species contain more tannin in nutrient poor environments (Hofland-Zijlstra and Berendse, 2009; Kouki and Manetas, 2002; Kraus et al., 2004; Northup et al., 1995). Our model resolves this apparent paradox (Fig. 3, middle panel). Tannins can be beneficial to plants growing in unfertile ecosystems because they allow the formation of recalcitrant nutrient pools less prone to leaching, which can counter-balance the negative effect of a slower mineralization (Schweitzer et al., 2008).

In the model of Barot et al. (2014), the nutrients present in dead organic matter could not be reabsorbed by plants directly, and had to be first decomposed into mineral nutrients, leading to the conclusion that plants must evolve fast mineralization strategies in unfertile ecosystems. Instead, in our model plants can have direct access to the recalcitrant pool and could thus benefit plants in unfertile ecosystems. In fact, it is more likely that fertile ecosystems *prevent* tannin evolution. In our model, plant-soil feedbacks initiated by tannins were inefficient at high fertility. Furthermore, fertile ecosystems favor fast growing species with high turn over rates (Endara and Coley, 2011). In this case we found that the selective pressure on tannins is weak (leftmost panel of Fig. 3).

### The importance of symbiosis with mycorrhizae

In our framework, tannin production and symbiotic associations with mycorrhizal fungi are two strongly interacting plant traits, are are thus likely to have co-evolved. Our model indeed predicts non-trivial coevolutionary dynamics that can lead to a coevolutionary equilibrium in which tannin production favors the association with mycorrhizal fungi and vice-versa (Fig 6). If plants benefit from a symbiotic association even when recalcitrant nutrient are absent, this coevolutionary equilibrium can be reached without any defensive role of tannins against herbivory (Fig 6, leftmost panel). If plants do not require symbiotic associations to absorb labile nutrients, a defensive role of tannins against herbivory, causing a build-up of recalcitrant nutrients unaccessible without symbiotic associations, can trigger the coevolutionary dynamics (Fig 6 rightmost panels).

These co-evolutionary dynamics rely on the ability of mycorrhizae to access nitrogen from tannin-protein complexes. This constitutes a central assumption in our work, which we now discuss.

Within functional groups of mycorrhizae, the mycorrhizal strategy may have evolved several times from free ancestors, as it is the case for ectomycorrhizae (Pellitier and Zak, 2018). Consequently, the different groups of mycorrhizae are not phylogenetically related and have distinct enzymatic capacity (Phillips et al., 2013). Their decomposer capacities are thus expected to depend on their type – ecto-,ericoid-or endomycorrhizae (for short: ECM, ERM, AMF) and of the clade within a given type.

That mycorrhizae can access recalcitrant complexes is still in debate for ECM (Lindahl and Tunlid, 2015; Pellitier and Zak, 2018), the dominant mycorrhizal type in temperate and boreal forests. The loss of genes associated with saprotrophic ability during the transition from a free to a mycorrhizal strategy, as well as certain experimental evidence (Bending and Read, 1996), suggest a low ability for complex organic matter decomposition. Simultaneously, an increasing body of litterature suggest the opposite, arguing that ECM selectively mine complex organic nitrogen (Averill and Hawkes, 2016; Shah et al., 2016; Trap et al., 2017). Madritch and Lindroth (2015) specifically demonstrated an higher N acquisition from high-tannin litter but without exploring the underlying mechanisms, in particular the potential implication of ECM fungi. Again, part of the controversy may arise from the fact that distinct ECM species might have distinct enzymatic abilities.

On the other hand, Wurzburger and Hendrick (2009), using 15 N-enriched protein-tannin complexes from leaf litter extracts, convincingly demonstrated the ability to acquire directly nitrogen from tannin-protein complexes for ERM, which often abound in heathland and peatlands, and are often very rich in tannins. AMF, which dominate in grasslands and tropical forests, are usually poor decomposers, and may therefore not be able to acquire nitrogen from tannin-protein complexes. However, as pointed by Hättenschwiler et al. (2010), this ability has so far not been tested for tropical AM fungi. In short, if our results should be relevant for ERM dominated ecosystems (e.g. heathland and peatlands), more studies are needed to clarify the decomposer ability of ECM and tropical AM fungi, and thus establish the precise level of generality of our results.

Incidentally, our study sheds light on the mycorrhizal-associated nutrient economy framework proposed by Phillips et al. (2013). This framework posits that the contrasting modes of nutrient acquisition of different mycorrhizal fungi can explain contrasting C and N cycling patterns observed in ecosystems dominated by ECM, ERM or AM plants. By modeling together the formation of recalcitrant N through tannin production and the ability of mycorrhizae to access it, our study refines this framework and places the patterns observed by Phillips et al. (2013) in an evolutionary perspective.

### Tannin evolution under a niche-construction perspective

The evolution of tannins through plant soil feedbacks can be understood as a case of niche construction, i.e. organisms’ ability to modify their environment for their own benefit, or for the benefit of related organisms which will occupy the improved envi-ronment after their death (Kylafis and Loreau, 2008; Lehmann, 2008; Odling-Smee et al., 2003). In our model tannins initiate an eco-evolutionary loop between organisms and their environments. The death of a tannin producing individual leaves a N-rich patch available for recruitment from neighbors that are likely to be of his kin. Individuals could inherit the environment transformed by their ancestors, a property termed ecological inheritance (Odling-Smee et al., 2003). As illustrated in Fig 5, this effect was substantial and positive in our model. Most evolutionary models consider the evolution of phenotypes whose fitness can be fully understood at the organismic level. In the case of tannin evolution, the fitness of this phenotype can be understood at the organismic level in the presence of herbivores, but not in the presence of the plant-soil feedbacks. The feedback involves plants, nutrient compartments, and mycorrhizae within the soil, and competition occurs between ecosystems having contrasted material cycles (Loreau, 1998, 2010). A whole ecosystem therefore persists through time via genetic and ecological inheritance. Within the framework of multi-level selection (Okasha, 2006), we therefore propose to interpret tannin evolution as the emergence of a new unit of selection at the ecosystem-level (Goodnight, 2000; Swenson et al., 2000).

### Conclusion

Our model shows that the protective role of tannins against herbivores reinforces their evolution, but is not a necessary condition. Plant-soil feedbacks can be strong enough to trigger alone tannin evolution, provided that the pool of recalcitrant nutrients complexed by tannins is, at least in part, reabsorbed by the mycorrhizae of the same plant, or by its neighboring relatives. Niche construction is here a central aspect of tannin evolution through plant-soil feedbacks, modulated by ecological conditions. Tannin evolution requires plants to have intermediate biomass turnover rates an inhabit relatively infertile ecosystems, thus highlighting the fact that niche construction is more effective in harsh environments. Such ecological conditions are generally fulfilled in ecosystems harboring tannin producing plants (Endara and Coley, 2011; McKey et al., 1978). Our model there-fore suggests that plant-soil feedbacks could have played a decisive role in the evolution of tannin production. More generally our work provides a detailed example of how plant traits, via their impact on a plant’s environment, can be selected via niche construction mechanisms implying the transformation of a whole local ecosystem.

## Acknowledgments

We thank Christiane Gallet for her helpful advices on an earlier version of this manuscript, and Matthieu Barbier for his help in programming the cellular automaton.

## Footnotes

1 *α_τ_* and *γ_τ_* are simple functions of *τ* and *α, γ_τ_* = (1 − *τ*)*γ*, and 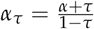.

2 The presence of *γB* in the photosynthesis term accounts for the fact that it is mostly N-compounds of biomass that perform photosynthesis.

## References

Aerts, R., and F. Chapin. 2000. The mineral nutrition of wild plants revisited: A reevaluation of processes and patterns. Pages 1–67 in Advances in Ecological Research, Vol 30, Advances in Ecological Research. Academic Press Inc, San Diego.

Allen, B., J. Gore, and M. A. Nowak. 2013. Spatial dilemmas of diffusible public goods. eLife 2:e01169.

Averill, C., and C. V. Hawkes. 2016. Ectomycorrhizal fungi slow soil carbon cycling. Ecology letters 19:937–947.

Barot, S., S. Bornhofen, N. Loeuille, N. Perveen, T. Shahzad, and S. Fontaine. 2014. Nutrient enrichment and local competition influence the evolution of plant mineralization strategy: a modelling approach. Journal of Ecology 102:357–366.

Bending, G., and D. Read. 1996. Nitrogen mobilization from protein-polyphenol complex by ericoid and ectomycorrhizal fungi. Soil Biology & Biochemistry 28:1603–1612.

Benoit, R., and R. Starkey. 1968. Enzyme inactivation as a factor in the inhibition of decomposition of organic matter by tannins. Soil Science 105:203–208.

Bergvall, U. A., and O. Leimar. 2005. Plant secondary compounds and the frequency of food types affect food choice by mammalian herbivores. Ecology 86:2450–2460.

Coq, S., J. Souquet, E. Meudec, V. Cheynier, and S. Hattenschwiler. 2010. Interspecific variation in leaf litter tannins drives decomposition in a tropical rain forest of French Guiana. Ecology 91:2080–2091.

Danger, M., T. Daufresne, F. Lucas, S. Pissard, and G. Lacroix. 2008. Does Liebig’s law of the minimum scale up from species to communities? Oikos 117:1741–1751.

Donohue, K., L. Dorn, C. Griffith, E. Kim, A. Aguilera, C. R. Polisetty, and J. Schmitt. 2005. Niche construction through germination cueing: life-history responses to timing of germination in Arabidopsis thaliana. Evolution 59:771–785.

Droop, M. R. 1968. The kinetics of uptake, growth and inhibition in Monochrysis lutheri. Journal of the Marine Biological Association of the United Kingdom 48:689–733.

Endara, M., and P. Coley. 2011. The resource availability hypothesis revisited: a meta-analysis. Functional Ecology 25:389–398.

Eshel, I. 1983. Evolutionary and continuous stability. Journal of theoretical Biology 103:99–111.

Feeny, P. 1970. Seasonal Changes in Oak Leaf Tannins and Nutrients as a Cause of Spring Feeding by Wintr Moth Caterpillars. Ecology 51:565–581.

Geritz, S. A., J. A. Metz, E. Kisdi, and G. Meszéna. 1997. Dynamics of adaptation and evolutionary branching. Physical Review Letters 78:2024.

Geritz, S. a. H., E. Kisdi, G. Meszéna, and J. a. J. Metz. 1998. Evolutionarily singular strategies and the adaptive growth and branching of the evolutionary tree. Evolutionary Ecology 12:35–57.

Goodnight, C. J. 2000. Heritability at the ecosystem level. Proceedings of the National Academy of Sciences 97:9365–9366.

Hättenschwiler, S., S. Coq, S. Barantal, and I. Handa. 2010. Leaf traits and decomposition in tropical forests: revisiting some commonly held views and towards a new hypothesis. New Phytologist.

Hättenschwiler, S., and P. Vitousek. 2000. The role of polyphenols in terrestrial ecosystem nutrient cycling. Trends in Ecology & Evolution 15:238–243.

Hofland-Zijlstra, J. D., and F. Berendse. 2009. The effect of nutrient supply and light intensity on tannins and mycorrhizal colonisation in Dutch heathland ecosystems. Plant Ecology 201:661–675.

Joanisse, G., R. Bradley, C. Preston, and A. Munson. 2007. Soil enzyme inhibition by condensed litter tannins may drive ecosystem structure and processes: the case of Kalmia angustifolia. New Phytologist 175:535–546.

Joanisse, G. D., R. L. Bradley, C. M. Preston, and G. D. Bending. 2009. Sequestration of soil nitrogen as tannin–protein complexes may improve the competitive ability of sheep laurel (Kalmia angustifolia) relative to black spruce (Picea mariana). New Phytologist 181:187–198.

Jordan, C., R. Todd, and G. Escalante. 1979. Nitrogen Conservation in a Tropical Rain Forest. Oecologia 39:123–128.

Kouki, M., and Y. Manetas. 2002. Resource availability affects differentially the levels of gallotannins and condensed tannins in Ceratonia siliqua. Biochemical Systematics and Ecology 30:631–639.

Kraus, T., R. Zasoski, and R. Dahlgren. 2004. Fertility and pH effects on polyphenol and condensed tannin concentrations in foliage and roots. Plant and Soil 262:95–109.

Kraus, T. E. C., R. A. Dahlgren, and R. J. Zasoski. 2003. Tannins in nutrient dynamics of forest ecosystems - a review. Plant and Soil 256:41–66.

Kylafis, G., and M. Loreau. 2008. Ecological and evolutionary consequences of niche construction for its agent. Ecology letters 11:1072–1081.

Kéfi, S., M. v. Baalen, M. Rietkerk, and M. Loreau. 2008. Evolution of local facilitation in arid ecosystems. The American Naturalist 172:E1–E17.

Laland, K. N., F. J. Odling-Smee, and M. W. Feldman. 1999. Evolutionary consequences of niche construction and their implications for ecology. Proceedings of the National Academy of Sciences 96:10242–10247.

Le Galliard, J.-F., R. Ferriére, and U. Dieckmann. 2003. The Adaptive Dynamics of Altruism in Spatially Heterogeneous Populations. Evolution 57:1–17.

Lehmann, L. 2008. The adaptive dynamics of niche constructing traits in spatially subdivided populations: evolving posthumous extended phenotypes. Evolution 62:549–566.

Leimar, O. 2009. Multidimensional convergence stability. Evolutionary Ecology Research 11:191–208.

Lemesle, V., and L. Mailleret. 2008. A mechanistic investigation of the algae growth “Droop” model. Acta biotheoretica 56:87–102.

Lewontin, R. C. 2001. The triple helix: Gene, organism, and environment. Harvard University Press.

Lindahl, B. D., and A. Tunlid. 2015. Ectomycorrhizal fungi–potential organic matter decomposers, yet not saprotrophs. New Phytologist 205:1443–1447.

Loreau, M. 1998. Ecosystem development explained by competition within and between material cycles. Proceedings of the Royal Society of London B: Biological Sciences 265:33–38.

Loreau, M. 2010. Evolution of ecosystems and ecosystem properties. Pages 225–259 in From Populations to Ecosystems, Theoretical Foundations for a New Ecological Synthesis. Princeton University Press.

Madritch, M. D., and R. L. Lindroth. 2015. Condensed tannins increase nitrogen recovery by trees following insect defoliation. New Phytologist.

McKey, D., G. Peter, C. Mbi, J. Gartlan, and T. Struhsaker. 1978. Phenolic Content of Vegetation in Two African Rain Forests: Ecological Implications. Science 202:61–64.

Mole, S. 1993. The systematic distribution of tannins in the leaves of angiosperms: A tool for ecological studies. Biochemical Systematics and Ecology 21:833–846.

Northup, R., Z. Yu, R. Dahlgren, and K. Vogt. 1995. Polyphenol Control of Nitrogen Release from Pine Litter. Nature 377:227–229.

Odling-Smee, F. J., K. N. Laland, and M. W. Feldman. 2003. Niche construction: the neglected process in evolution. 37. Princeton University Press.

Odum, E. P. 1971. Fundamentals of ecology. 3rd ed. Saunders, Philadelphia.

Okasha, S. 2006. Evolution and the Levels of Selection, vol. 16. Clarendon Press Oxford.

Ordoñez, J. C., P. M. Van Bodegom, J.-P. M. Witte, I. J. Wright, P. B. Reich, and R. Aerts. 2009. A global study of relationships between leaf traits, climate and soil measures of nutrient fertility. Global Ecology and Biogeography 18:137–149.

Pellitier, P. T., and D. R. Zak. 2018. Ectomycorrhizal fungi and the enzymatic liberation of nitrogen from soil organic matter: why evolutionary history matters. New Phytologist 217:68–73.

Phillips, R. P., E. Brzostek, and M. G. Midgley. 2013. The mycorrhizal-associated nutrient economy: a new framework for predicting carbon–nutrient couplings in temperate forests. New Phytologist 199:41–51.

Pritsch, K., and J. Garbaye. 2011. Enzyme secretion by ECM fungi and exploitation of mineral nutrients from soil organic matter. Annals of Forest Science 68:25–32.

Read, D., and J. Perez-Moreno. 2003. Mycorrhizas and nutrient cycling in ecosystems - a journey towards relevance? New Phytologist 157:475–492.

Schweitzer, J., J. Bailey, B. Rehill, G. Martinsen, S. Hart, R. Lindroth, P. Keim, and T. Whitham. 2004. Genetically based trait in a dominant tree affects ecosystem processes. Ecology Letters 7:127–134.

Schweitzer, J., I. Juric, T. Voorde, K. Clay, W. Putten, and J. Bailey. 2014. Are there evolutionary consequences of plant–soil feedbacks along soil gradients? Functional Ecology 28:55–64.

Schweitzer, J. A., M. D. Madritch, J. K. Bailey, C. J. LeRoy, D. G. Fischer, B. J. Rehill, R. L. Lindroth, A. E. Hagerman, S. C. Wooley, and S. C. Hart. 2008. From genes to ecosystems: the genetic basis of condensed tannins and their role in nutrient regulation in a Populus model system. Ecosystems 11:1005–1020.

Schwilk, D. W. 2003. Flammability is a niche construction trait: canopy architecture affects fire intensity. The American Naturalist 162:725–733.

Shah, F., C. Nicolás, J. Bentzer, M. Ellström, M. Smits, F. Rineau, B. Canbäck, D. Floudas, R. Carleer, and G. Lackner. 2016. Ectomycorrhizal fungi decompose soil organic matter using oxidative mechanisms adapted from saprotrophic ancestors. New Phytologist 209:1705–1719.

Swenson, W., D. S. Wilson, and R. Elias. 2000. Artificial ecosystem selection. Proceedings of the National Academy of Sciences 97:9110–9114.

Trap, J., M. Akpa-Vinceslas, P. Margerie, S. Boudsocq, F. Richard, T. Decaëns, and M. Aubert. 2017. Slow decomposition of leaf litter from mature Fagus sylvatica trees promotes offspring nitrogen acquisition by interacting with ectomycorrhizal fungi. Journal of Ecology 105:528–539.

van Baalen, M., and D. A. Rand. 1998. The unit of selection in viscous populations and the evolution of altruism. Journal of Theoretical Biology 193:631–648.

Vitousek, P. 1982. Nutrient cycling and nutrient use efficiency. The American Naturalist 119:553–572.

Whitham, T. G., J. K. Bailey, J. A. Schweitzer, S. M. Shuster, R. K. Bangert, C. J. LeRoy, E. V. Lonsdorf, G. J. Allan, S. P. DiFazio, and B. M. Potts. 2006. A framework for community and ecosystem genetics: from genes to ecosystems. Nature Reviews Genetics 7:510–523.

Wilson, D. S. 1980. The natural selection of populations and communities. Benjamin/Cummings Pub. Co.

Wurzburger, N., and R. Hendrick. 2009. Plant litter chemistry and mycorrhizal roots promote a nitrogen feedback in a temperate forest. Journal of Ecology 97:528–536.

